# Reticulocytes in donor RBC units enhance RBC alloimmunization

**DOI:** 10.1101/2023.01.25.525560

**Authors:** Tiffany A. Thomas, Annie Qiu, Christopher Y. Kim, Dominique E. Gordy, Anabel Miller, Maria Tredicine, Monika Dzieciatkowska, Flavia Dei Zotti, Eldad A. Hod, Angelo D’Alessandro, James C. Zimring, Steven L. Spitalnik, Krystalyn E. Hudson

## Abstract

Although red blood cell (RBC) transfusions save lives, some patients develop clinically-significant alloantibodies against donor blood group antigens, which then have adverse effects in multiple clinical settings. Few effective measures exist to prevent RBC alloimmunization and/or eliminate alloantibodies in sensitized patients. Donor-related factors may influence alloimmunization; thus, there is an unmet clinical need to identify which RBC units are immunogenic. Repeat volunteer blood donors and donors on iron supplements have elevated reticulocyte counts compared to healthy non-donors. Early reticulocytes retain mitochondria and other components, which may act as danger signals in immune responses. Herein, we tested whether reticulocytes in donor RBC units could enhance RBC alloimmunization. Using a murine model, we demonstrate that transfusing donor RBC units with increased reticulocyte frequencies dose-dependently increase RBC alloimmunization rates and alloantibody levels. Transfusing reticulocyte-rich RBC units was associated with increased RBC clearance from the circulation and a robust proinflammatory cytokine response. As compared to previously reported post-transfusion RBC consumption patterns, erythrophagocytosis from reticulocyte-rich units was increasingly performed by splenic B cells. These data suggest that reticulocytes in a donated RBC unit impact the quality of blood transfused, are targeted to a distinct compartment, and may be an underappreciated risk factor for RBC alloimmunization.

## Introduction

Red blood cells (RBCs) are transfused to correct anemia from bone marrow failure, hemolytic anemia, and hemorrhage. Although RBC transfusions can be lifesaving, some patients develop alloantibodies against donor RBC blood group antigens. These alloantibodies can be clinically significant, inducing hemolytic transfusion reactions and hyperhemolysis, raising transplantation barriers, and delaying transfusion by making it difficult to find compatible blood for transfusion(1). Once alloantibodies arise, patient care becomes more challenging. Currently, there are few strategies to prevent RBC alloimmunization besides antigen matching or antigen avoidance, and few treatments are available (with limited efficacy) once alloimmunization occurs(1).

As opposed to pharmaceuticals, each unit of RBCs is unique as it comes from a distinct individual, undergoes a number of different manufacturing steps, and then is followed by a variable duration of refrigerator storage up to 42 days before transfusion(2, 3). Increasing evidence suggests that the risk of alloimmunization following an RBC transfusion is influenced by donor factors (e.g., blood group antigen density), component factors (e.g., storage duration) and host factors (e.g., blood group antigen-negativity, HLA type, inflammation)(4). Identifying new risk factors that can prevent and/or reduce risk of future RBC alloimmunization events is a priority, given the current lack of available treatments for sensitized patients. In both humans and mice, RBC alloimmunization rates increase in transfusion recipients with viral infections or viral-like inflammation(4, 5). Viruses induce robust type I interferon (IFN-α/β) production; in mouse models of RBC alloimmunization, blocking IFN receptors prevents alloantibody production, whereas infusing IFN-α promotes alloantibodies(6, 7). Mechanistically, viral-like inflammation stimulates splenic antigen presenting cells (APCs) and alters the consumption of transfused RBCs towards immunogenic dendritic cell (DC) subsets and away from red pulp macrophages(8, 9).

Intriguingly, patients at highest risk for RBC alloimmunization (e.g., those with sickle cell disease (SCD)) have elevated type I IFN levels(10-12). Although the mechanisms underlying type I IFN production are unknown, we and others recently published that mitochondria in human RBCs stimulate type I IFN production by immune cells(13, 14). Although mitochondrial retention in mature RBCs is abnormal, most early reticulocytes have identifiable mitochondria. Some transfusion recipients and repeat volunteer blood donors have elevated reticulocyte frequencies(15, 16). Moreover, because compensatory reticulocytosis lasts longer than the interval between donations, RBC units from some repeat volunteer blood donors have reticulocyte counts above the reference range(15). Of note, stress-induced reticulocytes produced in response to anemia (e.g., phlebotomy, hemorrhage) are larger, less deformable, and contain higher numbers of mitochondria(17-23). Thus, we hypothesized that reticulocytes in RBC donor units may enhance RBC alloimmunization.

Using a murine model of RBC alloimmunization, we test the hypothesis that elevated reticulocyte counts in RBC donor units enhance RBC alloantibody production. We demonstrate a direct correlation between reticulocyte counts in an RBC donor unit and RBC alloantibody levels produced in transfusion recipients. Compared to controls, reticulocyte-rich RBC donor units were enriched in mitochondrial proteins and metabolites and, upon transfusion, elicited proinflammatory cytokines. Mechanistically, RBCs from reticulocyte-rich units had lower 24-hour post-transfusion recovery (i.e., were cleared more robustly in the first 24 hours after transfusion) and induced significantly higher RBC consumption by splenic APCs. Multiple B cell subsets consumed most of the reticulocytes, representing a departure from normal post-transfusion RBC consumption patterns. The effect of higher reticulocyte frequencies was durable because transfusion of short- (i.e., 1 day) and medium- (i.e., 6 day) stored RBC donor units similarly induced alloantibodies. Lastly, we show that reticulocyte-mediated enhanced alloimmunization is not Toll-Like Receptor (TLR)-4 dependent. Together, these data demonstrate that reticulocytes in RBC donor units can enhance alloimmune responses.

## Materials and Methods

### Mice

HOD mice(24), expressing an RBC-specific triple fusion protein consisting of hen egg lysozyme, ovalbumin, and human blood group molecule Duffy, were bred in the Columbia University vivarium. C57BL/6 (B6, strain #027) mice were purchased from Charles River Laboratories. C57BL/6-Tg(UBC-GFP)30Scha/J (GFP; strain #004353) and B6(Cg)-Tlr4^tm1.2Karp^/J (TLR4^-/-^, strain #029015) mice were purchased from The Jackson Laboratory. All mice were on a 12:12 light/dark cycle and maintained on a chow diet (ad libitum) unless otherwise specified. All murine experiments were approved by Columbia University’s Institutional Animal Care and Use (IACUC) committee.

### Reticulocyte induction and blood bank preparation

#### Reticulocyte induction

Reticulocytosis was induced with phenylhydrazine (PHZ), or by iron deficiency followed by acute iron repletion. **PHZ:** Intravascular hemolysis was induced with 2 intraperitoneal (i.p.) injections of PHZ (Sigma) 50mg/kg/day separated by 24 hours or saline control. **Iron:** Weanling (3-week-old) mice were placed on iron deficient (Harlan Teklad TD.110592) or replete (Harlan Teklad TD.110593) diets for 4 weeks, followed by reticulocytosis induction by intraperitoneal injection of 5mg of iron dextran (0.1mg of 50mg/mL solution; Allergan NDC 00230608-10) and switching to iron replete chow(25). <u>Blood bank preparation.</u> Whole blood was collected by cardiac puncture into 14% CPDA-1 four days after PHZ or iron dextran injection, pooled, filter leukoreduced (Acrodisc WBC syringe filter, Pall), packed to 60% hematocrit, and stored for 1 or 6 days at 4°C in 1.5mL microcentrifuge tubes. <u>Reticulocyte quantification.</u> 1μL of blood was incubated with 2.5nM of MitoTracker Deep Red FM (ThermoFisher) for 30 min at 37°C. After washing with FACS buffer, samples were stained with antibodies against CD45, CD41, and CD71 for 30 min at 4°C (**Supplemental Table 1**)(13). After washing with FACS buffer, cells were analyzed using an Attune NxT flow cytometer (ThermoFisher). CD41+ platelets and CD45+ white blood cells were excluded and RBCs were identified using a 405nm filter (i.e., No wash no lyse filter, ThermoFisher), which leverages the properties of hemoglobin. RBCs were distinguished by CD71 expression: CD71+ reticulocytes and CD71-mature RBCs. Data were analyzed with FlowJo software (BD Biosciences).

### RBC transfusion, post-transfusion recovery (PTR), and RBC alloantibody detection

On the morning of PTR experiments, a leukoreduced RBC unit (60% hematocrit) was prepared from mice whose RBCs were biotinylated *in vivo* (1mg/300μL in PBS injected into the retro-orbital sinus)(26). Stored RBC units were spiked 1:5 with fresh biotinylated RBC units as a loading control. Aliquots (200μL) from these spiked RBC units were transfused into recipients via the retro-orbital venous sinus. Blood samples were collected via tail puncture (1μL in 1mL PBS) at 5 minutes, 120 minutes, and 24 hours after transfusion. PTR was measured as the ratio of biotin-positive RBCs to GFP-positive RBCs circulating 24 hours-post transfusion relative to the same ratio in the spiked blood unit. Sera were collected from recipients and total HOD alloantibodies (IgM + IgG + IgA) were quantified by flow crossmatch(27).

### Hematocrit and hemoglobin

Whole blood was collected from the submandibular vein into microcentrifuge tubes containing EDTA. Hematocrit was determined following micro-hematocrit capillary tube centrifugation. Hemoglobin was calculated spectrophotometrically (540nm) using Drabkin’s reagent.

### Cytokine production and splenic erythrophagocytosis

Recipients were transfused with 6-day stored PHZ, stored control, or freshly collected GFP RBC units (350μL). <u>Cytokines.</u> Plasma collected 2 hours post-transfusion was diluted 1:1 in duplicate. Cytokines were quantified by a fluorescence-encoded bead-based multiplex assay (LEGENDplex Mouse Anti-Virus Response Panel, BioLegend), per manufacturer’s instructions. <u>Erythrophagocytosis.</u> Spleens were excised into complete RPMI, finely chopped, and filtered, as previously described(9, 27, 28). Single cells were washed with FACS buffer, RBCs were lysed, and the remaining cells stained with antibodies in FACS buffer to delineate leukocyte subsets (**Supplemental Table 1**)(8, 9, 28, 29). Results were acquired on an Attune NxT flow cytometer (ThermoFisher) and data analyzed with FlowJo software. Cell images were acquired on an Amnis ImageStreamX MkII cytometer (Luminex); 20,000 CD19+ B cells per file were recorded using INSPIRE software at 60x magnification. Analysis was performed using IDEAS 6.3 software.

### Ultra-High-Pressure Liquid Chromatography (UHPLC)–Mass Spectrometry (MS) Metabolomics

Metabolomics analyses were performed using a Vanquish UHPLC coupled online to a Q Exactive mass spectrometer (ThermoFisher, Bremen, Germany), as previously described(30). Data were analyzed using Maven (Princeton University) and Compound Discoverer 2.1 (ThermoFisher). Graphs and statistical analyses were prepared with MetaboAnalyst 5.0(31).

### Protein Digestion

Protein pellets from RBC units were digested as previously described(32).

### Nano Ultra-High-Pressure Liquid Chromatography–Tandem Mass Spectrometry Proteomics

Sample processing and data collection were performed as previously(32).

### Statistical Analysis

A repeated measures 2-way ANOVA with Sidak’s multiple comparisons test was utilized for analysis of alloantibody production over time. For comparison of 3 or more groups, a one-way ANOVA with Tukey’s multiple comparisons post-test was utilized. An unpaired t-test was used to compare 2 groups; p<0.05 was considered significant. Analyses were performed using GraphPad Prism, version 9.4.1.

## Results

### Transfusion of stored reticulocyte-rich RBC units enhances RBC alloimmunization

To test whether reticulocytes in RBC units modulate alloimmunization, RBC alloantibody production was evaluated in mice transfused with allogeneic RBC units containing elevated reticulocyte counts (**experimental design in Fig. 1A**). To generate RBC units enriched in reticulocytes, reticulocytosis was induced by phenylhydrazine (PHZ); PHZ damages RBCs, leading to hemolytic anemia followed by stress-induced reticulocytosis(33). A PHZ dose titration revealed that CD71+ reticulocyte frequency peaked in peripheral blood 3-4 days post treatment, with an optimal dose of 50mg/kg (**Supplemental Fig. 1**). Thus, HOD RBC donor mice, which express an RBC-specific HOD alloantigen, were treated with PHZ or saline control to generate RBC units. Compared to control, reticulocyte-rich RBC units had higher reticulocyte levels (**2% vs 50%, p<0.0001; Fig. 1B**) and of the reticulocytes, an increased frequency of mitochondria-positivity (**51% vs 64%, p<0.0001; Fig. 1C**), as determined using MitoTracker dye. Allogeneic HOD RBC units were refrigerator-stored 6 days before transfusion into wild-type B6 animals; sera were collected weekly and assessed for anti-HOD alloantibodies by flow crossmatch. Transfusion of stored reticulocyte-rich (referred to as “reticulocytes”) RBC units induced significantly higher HOD alloantibody levels, compared to stored saline control RBCs (“control”) (**p<0.001; Fig. 1D**), with an >800-fold difference in RBC alloantibody levels in recipient mice transfused with reticulocyte-rich RBC units, compared to control (**Fig. 1E**). Additionally, all recipients transfused with reticulocyte-rich RBC units had detectable alloantibodies, compared to some animals in the control transfusion group that did not respond. To test whether the enhanced RBC alloimmunization was due to the method of inducing reticulocytosis, RBC units with higher reticulocyte frequencies were also generated using an iron-deficient/iron dextran injection mouse model. As with PHZ-induced reticulocytosis, transfusing reticulocyte-rich RBC units from iron-deficient/iron dextran injection donors induced significantly higher anti-HOD alloantibody levels, compared to controls (**Supplemental Fig. 2**). Together, these data demonstrate that transfusing reticulocyte-rich RBC units induces higher RBC alloimmunization rates and alloantibody levels.

**Figure 1:**
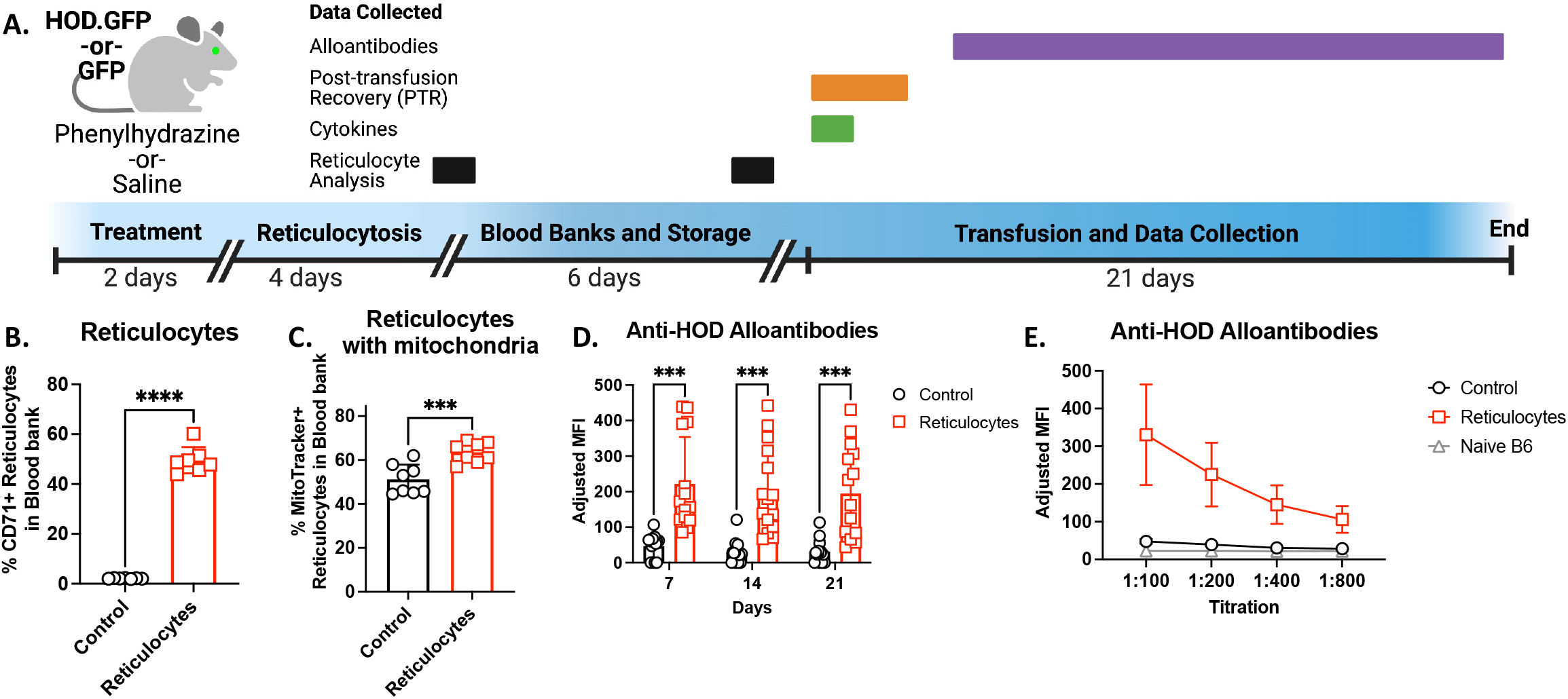
Transfusion of refrigerator-stored reticulocyte-rich RBC units enhanced RBC alloimmunization. (A) General experimental timeline. HOD mice received 2 i.p. injections of phenylhydrazine (PHZ, 50mg/kg, referred to as “Reticulocytes”)) or saline control (referred to as “Control”) spaced one day apart. Four days after treatment, whole blood was collected by cardiac puncture into 14% CPDA-1, leukoreduced, packed to a 60% hematocrit, and stored for 6 days at 4°C before transfusion. For some experiments, on the day of transfusion, fresh RBCs were collected from HOD mice as an additional control. (B) The percent reticulocytes and (C) the percent of mitochondria-positive reticulocytes generated following PHZ and saline treatment. (D) Post-transfusion sera were analyzed for total anti-HOD RBC alloantibodies by flow crossmatch. (E) Sera collected at 14 days post transfusion were titrated. Naïve B6 sera were run in triplicate to establish the background antibody level. Data are cumulative from 3 independent experiments with 5 mice/group. Data with 2 groups were analyzed with an unpaired T-test whereas 3 groups were analyzed by a repeated measures one-way ANOVA with Sidak’s multiple comparisons test and ***p<0.001, ****p<0.0001. Timeline figure created with BioRender.

### Transfusion of stored reticulocyte-rich RBC units enhances proinflammatory cytokine production

To test whether transfusion of reticulocyte-rich RBC units elicited an inflammatory response, plasma was collected two hours post-transfusion and assayed for cytokines. Transfusion of reticulocyte-rich RBC units led to significant increases in MCP-1, CXCL1, CXCL10, and IFN-γ levels, compared to control (**Fig. 2A**). No significant differences were noted in plasma IL-6, CCL5, IL-12, TNF-α, GM-CSF, IFN-α, IFN-β, IL-10, or IL-1β levels (**data not shown**). Together, these data suggest that transfusion of reticulocytes elicited a strong, albeit selective, inflammatory response, characterized by significant chemokine production.

**Figure 2:**
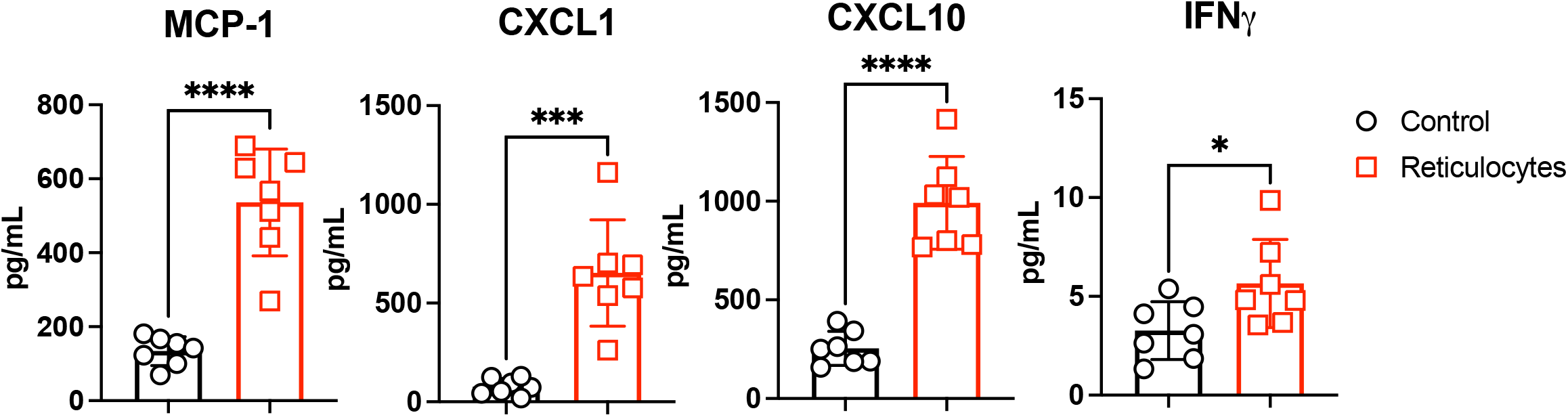
Induction of inflammatory cytokines after transfusing refrigerator-stored reticulocyte-rich RBC units. Reticulocyte-rich and control RBC units were generated (as in Figure 1) using whole blood collected from GFP animals. RBC units were stored for 6 days and transfused into wild-type B6 animals. Plasma was collected 2 hours post-transfusion and analyzed for cytokines. Data shown are cumulative of 2 independent experiments with 3-4 mice/group. Data were analyzed with an unpaired t-test; ****p<0.0001, ***p<0.001, *p<0.05.

### Reticulocyte-rich RBC units have elevated levels of mitochondrial metabolites and proteins

Untargeted and targeted metabolomics analyses using UHPLC-MS were performed on 6-day stored reticulocyte-rich and control RBC units; detailed results are provided in **Supplemental Table 2**. The metabolic phenotypes of reticulocyte-rich blood differed substantially from controls. Hierarchical clustering analysis (**Fig. 3A**) highlighted significant associations of reticulocytes with metabolic pattern differences in lipid (fatty acid, phospholipid, and sphingolipid), amino acid, and purine metabolism. Volcano plots (**Fig. 3B**) from targeted metabolomic analysis highlighted the top metabolites that increased (red) or decreased (blue) in the reticulocyte-rich RBC units as compared to controls. Pathway analysis of untargeted metabolomics data (**Fig. 3C**) revealed altered regulation of metabolic pathways suggestive of active mitochondrial function (i.e., beta oxidation, TCA cycle, urea cycle, malate-aspartate shuttle, and glycerol-3-phosphate shuttle) and mitochondrial synthesis (cardiolipin synthesis).

**Figure 3:**
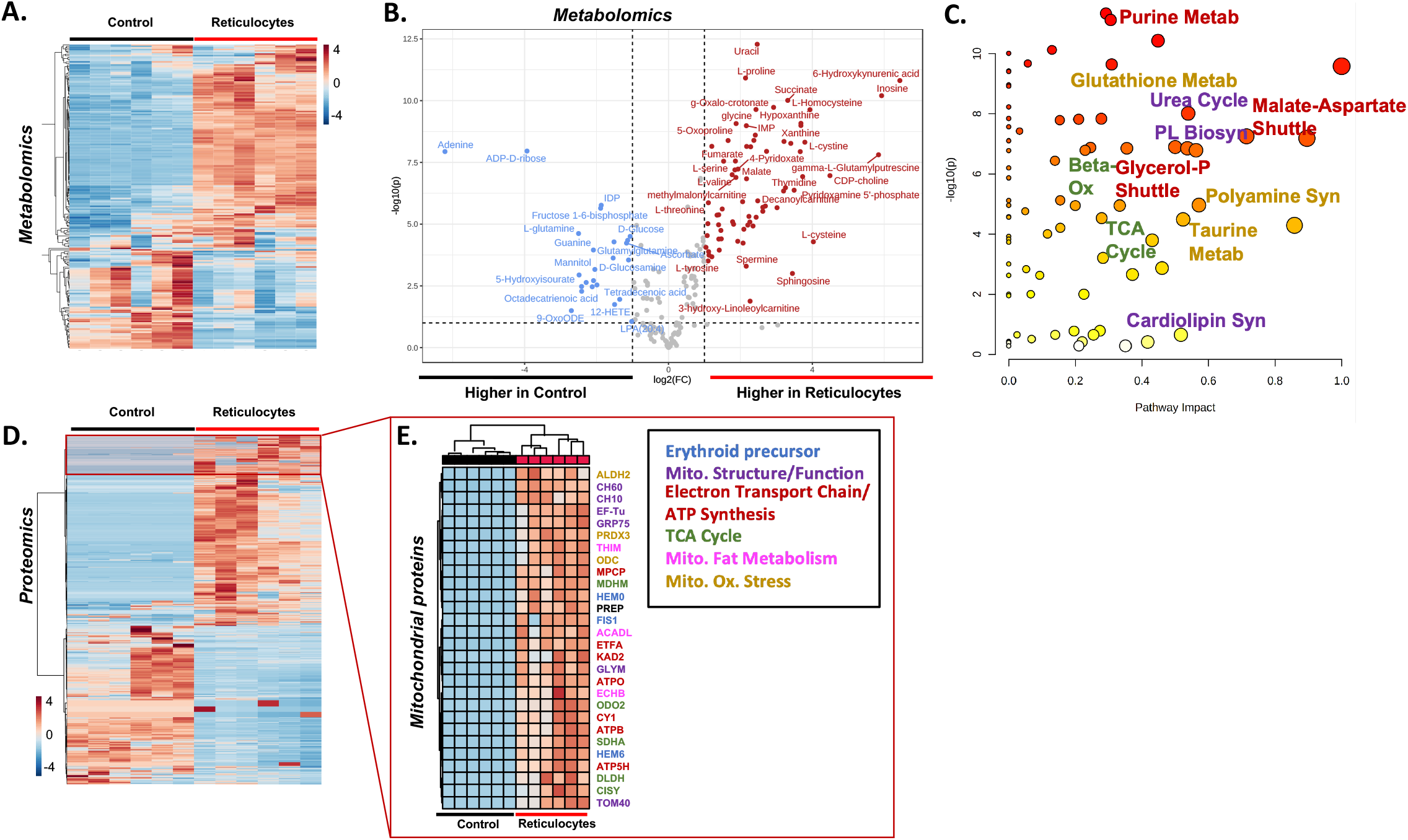
Omics analysis revealed mitochondrial metabolites and proteins in reticulocyte-rich RBC units. Aliquots of 6-day stored reticulocyte and control RBC units were subjected to omics analysis. (A) Hierarchical clustering analysis and (B) volcano plot highlighting significant metabolic changes between control and reticulocyte-rich RBC units. (C) Pathway analysis of untargeted metabolomics data. (D) Hierarchical clustering of significantly different proteins between control and reticulocyte-rich RBC units. In (E), zoom in on a subset of mitochondrial proteins (by gene ontology of cell localization) whose levels are higher in reticulocyte-rich RBC units, compared to controls.

Proteomics analyses were also performed on 6-day stored reticulocyte-rich and control RBC units. Hierarchical clustering analysis (**Fig. 3D**) highlighted significant associations of reticulocytes with proteins that were labeled of mitochondrial origin upon gene ontology enrichment analysis for cell localization, with the most differentially expressed proteins involved in mitochondrial fatty acid metabolism, the TCA cycle, electron transport chain, ATP synthesis, mitochondrial structure, and mitochondrial oxidative stress (**Fig. 3E**).

### Stored reticulocyte-rich RBC units have reduced post-transfusion recovery and are preferentially ingested by splenic B cells

Based on the enhanced cytokine production by, and the detectable mitochondrial byproducts in, reticulocyte-rich RBC units, we hypothesized that clearance of transfused RBCs from the circulation would be accelerated. To test this, recipients were transfused with reticulocyte-rich or control RBC units and PTR (i.e., the percentage of transfused RBCs remaining in the circulation for 24 hours) determined. Reticulocyte-rich RBCs had significantly reduced PTR at 24 hours, compared to control (**p<0.0001; Fig. 4A**,). To elucidate which cells were responsible for RBC clearance, reticulocyte-rich and control RBC units were generated using GFP mouse donors. Recipient B6 mice were then transfused with stored reticulocyte-rich, stored control, or freshly collected GFP RBC units and spleens harvested from recipients 2 hours later. Total splenocyte numbers were calculated and cells were stained with antibodies to identify antigen presenting cell (APC) subsets. For analysis, T cells and RBCs (or cells with RBCs attached to their surface) were excluded from total live splenocytes by excluding those positive for Thy1.2 and/or Ter119; GFP fluorescence provided an indirect measure of RBC consumption(8, 9, 28). Enumerating leukocytes revealed a significant decrease in total cell numbers and in APC subsets, including F4/80+ macrophages, CD8+ DCs, and CD19+ B cells, in recipients transfused with reticulocyte-rich RBC units, compared to stored and fresh control RBC units (**Fig. 4B-C**). No significant numerical differences in CD11b+ DCs were noted. Evaluating leukocytes for GFP fluorescence showed a significant increase in the frequency of GFP+ cells, suggesting that transfused reticulocyte-rich RBC units were being consumed by more leukocytes as compared to controls (**p<0.0001; Fig. 4D**). Individual APC subsets were interrogated for GFP fluorescence to determine which participated in erythrophagocytosis. As previously reported(34), transfusing stored control RBC units induced increased RBC consumption by macrophages, CD8+ DCs, and CD11b+ DCs, compared to fresh GFP control RBCs (**Fig. 4E**). However, as compared to stored control RBC units, consumption of RBCs from reticulocyte-rich units was significantly increased in CD8+ and CD11b+ DCs; nonetheless, consumption by macrophages was not enhanced and was similar to fresh RBC controls. Unexpectedly, B cells markedly consumed RBCs from transfused reticulocyte-rich units, compared to stored and fresh controls. These data demonstrate that transfusion of reticulocyte-rich RBC units leads to significant changes in splenocyte cell subset composition and RBC consumption patterns. The most striking finding was that >20% of B cells had detectable GFP fluorescence, suggesting that a unique RBC consumption pattern may be associated with stored reticulocyte clearance.

**Figure 4:**
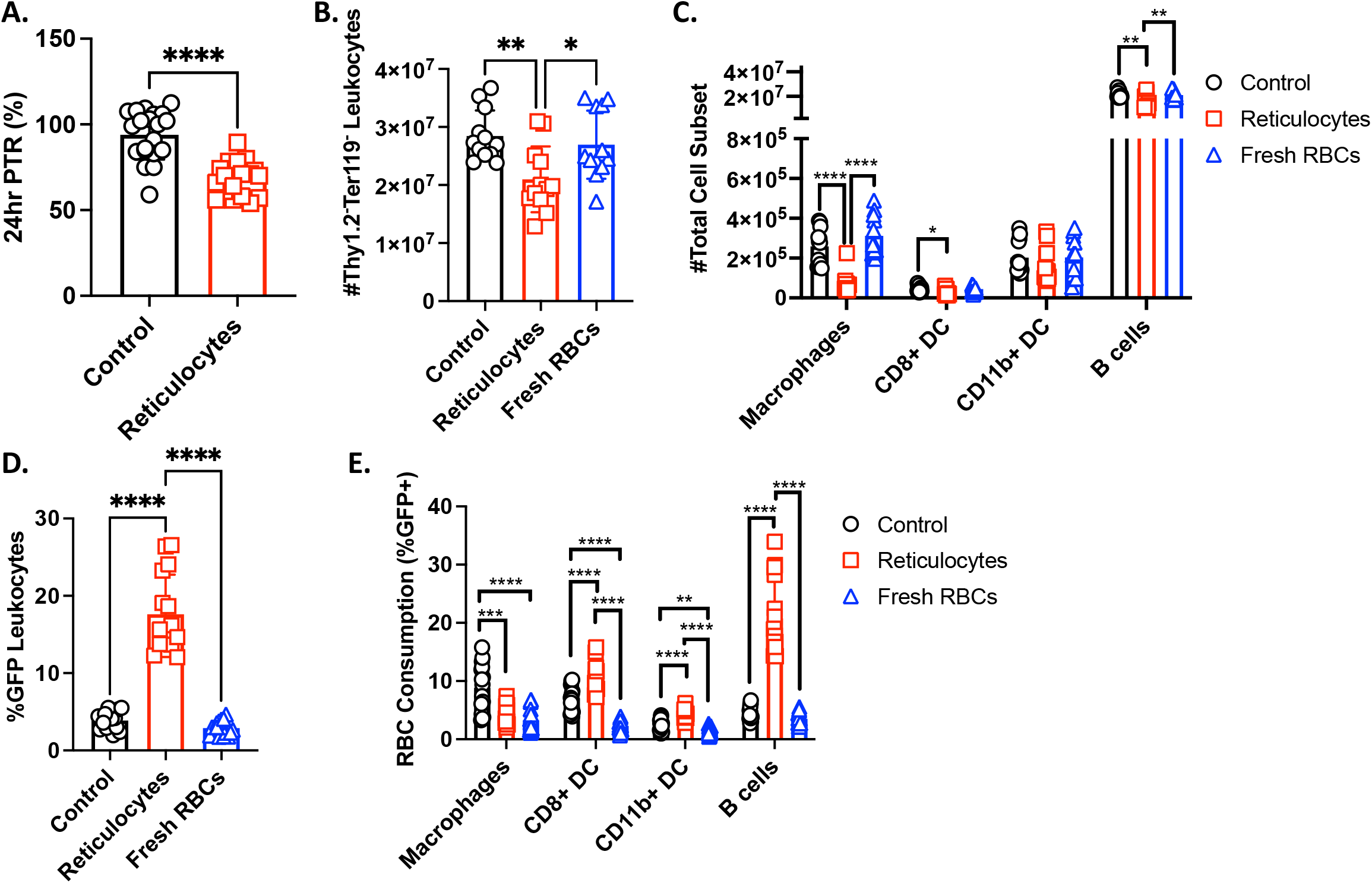
Transfusion with refrigerator-stored reticulocyte-rich RBC units led to increased RBC consumption. To determine post-transfusion recovery (PTR), reticulocyte-rich and control RBC units were generated from UBC.GFP mice. After 6 days of storage, RBC units were spiked with freshly-biotinylated freshly-obtained RBCs (as a tracer) and 350μL were transfused into wild-type B6 recipients. (A) Whole blood was sampled 24 hours later and the frequency of GFP+ RBCs was normalized to biotinylated RBCs to determine PTR. To evaluate cellular sites of RBC consumption, B6 recipients were transfused with 350μL of freshly collected GFP+ RBCs or 6-day stored RBC units (i.e., reticulocyte or control). Spleens were collected 2 hours post-transfusion and splenocytes stained with antibodies to delineate APC subsets. GFP signal level was used as an indirect measure of erythrophagocytosis. The total number of cells per spleen were calculated for (B) Thy1.2-Ter119-leukocytes and for (C) individual APC subsets. The frequencies of GFP+ (D) Thy1.2-Ter119-leukocytes and (E) individual APC subsets were determined. The following phenotypes were used to delineate APC subsets: macrophages: CD11c^-/lo^CD11b^-/lo^F4/80^+^; CD8^+^ DCs: CD11c^hi^CD8^+^CD11b^-^; CD11b^+^ DCs: CD11c^hi^CD11b^+^CD8^-^; and B cells: CD19^+^. Data are aggregated from at least 3 independent experiments with 3-4 mice/group. Data were analyzed with an unpaired t test for 2 groups or one way ANOVA with Tukey’s multiple comparisons test; ****p<0.0001, ***p<0.001, **p<0.01, *p<0.05

To elucidate which splenic B cell subsets were involved in clearing reticulocyte-rich RBC units, B cells were gated into follicular, marginal zone, and B1 B cell subsets (gating strategy shown in **Supplemental Fig 3**) (29). Low levels of erythrophagocytosis by B cell subsets were observed upon transfusion with stored or fresh control RBC units (**Fig. 5A**). In contrast, there was a marked increase in GFP+ B cells following transfusion with reticulocyte-rich units. The most prominent signal was with innate B1 B cells, with ∼45% positive for GFP. To confirm whether B cells actually internalized RBCs, samples were analyzed with an imaging cytometer. With reticulocyte-rich samples, three GFP expression patterns emerged: diffuse GFP signal within CD19+ staining, punctate GFP signal along the cell membrane, and clusters of GFP signal within the B cell (**Fig. 5B, left; Supplemental Fig. 4A-B**). In contrast, in control samples, only punctate GFP signal along the B cell membrane was detectable, along with a few cells that had RBC particulates attached to the B cell surface (**Fig. 5B, right**). In samples from the fresh control RBC transfusion group, very few GFP+ B cells were detected and most positive events reflected RBC particulates attached outside the B cell, as indicated by brightfield images (**Supplemental Fig. 4C**); this was consistent with the frequency of cells with GFP signal inside the cell membrane (**Fig. 5C**).

**Figure 5:**
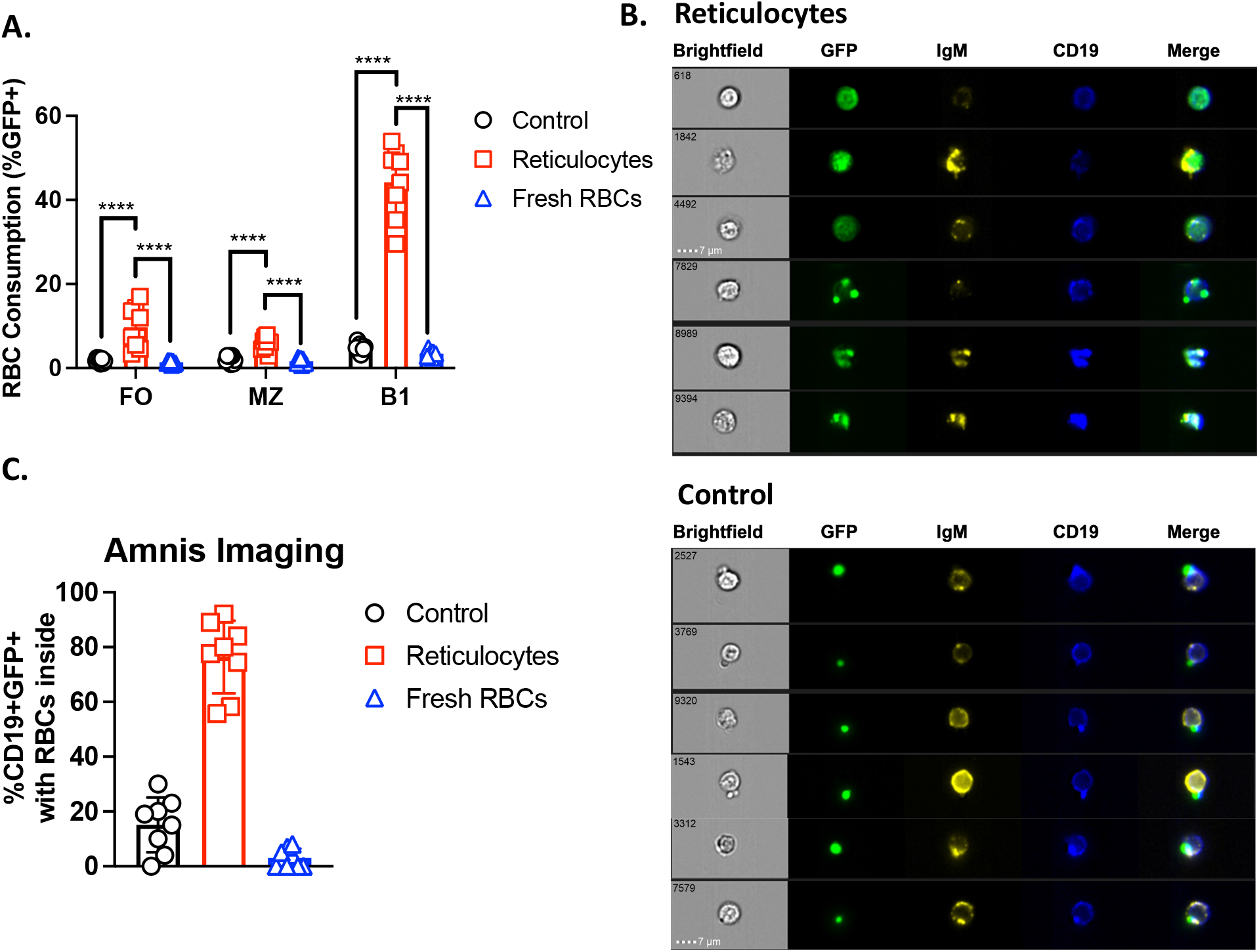
Multiple B cell subsets consume stored reticulocytes. Recipients were transfused with 350μL of freshly collected GFP+ RBCs or 6-day stored RBCs collected from PHZ or saline-treated mice. Spleens were collected 2 hours post-transfusion and splenocytes stained with antibodies to delineate APC subsets. GFP signal level was used as an indirect measure of erythrophagocytosis. (A) Total frequency of GFP+ cells for each B cell subset was calculated by flow cytometry. (B) Imaging analysis of B cells from mice transfused with refrigerator-stored reticulocyte-rich (top) or stored saline (bottom) RBC units. Cell #4492 demonstrates RBC internalization by a B cell whereas #2527 represents an RBC adhered to the surface of a B cell. (C) Percentage of CD19+GFP+ cells with GFP+ RBCs located inside their cell membrane. The phenotypes used to delineate CD19+ B cell subsets were follicular (FO; B220^+^IgD^+^IgM^lo^CD23^+^CD93^-^), marginal zone (MZ; B220^+^IgD^+^IgM^+^CD23^-^CD93^-^), and B1 (B220^lo/-^CD43^+^). To exclude RBCs attached to the external B cell surface, Ter119+ cells were excluded from parent gates. Data are cumulative from 2 experiments with n=3-4 mice/group. Data were analyzed with one-way ANOVA with Tukey’s multiple comparisons test; ****p<0.0001

### RBC alloimmunization is enhanced, even after shorter storage and fewer reticulocytes

Because refrigerator storage adversely affects RBC quality and because reticulocyte counts are much higher in reticulocyte-rich RBC units generated from mice, as compared to volunteer human donors, we generated mouse RBC units with defined reticulocyte frequencies. HOD RBC units were refrigerator-stored for 1 day (equivalent to ∼3 days of storage for human RBC units) and then transfused into recipients. Transfusing RBC units with as few as 5% reticulocytes led to significant alloantibody production, as compared to control (i.e., <2% reticulocytes) (**Fig. 6**). There was a significant direct correlation between reticulocyte frequency in the RBC donor unit and alloantibody production; this relationship was significant even at low reticulocyte frequencies. Lastly, to test whether B cell subsets consumed RBCs from 1 day stored reticulocyte-rich units, GFP fluorescence of follicular, marginal zone, and B1 B cell subsets was assessed; significant RBC consumption was observed in each of these B cell subsets following reticulocyte-rich RBC transfusions, compared to controls (**Fig. 6B**). Thus, B cell consumption patterns were similar between 1 and 6 day refrigerator-stored reticulocyte-rich RBC units. These findings were visually confirmed by imaging cytometry (**Fig. 6C, Supplemental Fig. 5**). Collectively, these data show that even low reticulocyte counts make RBC units more immunogenic, and that prolonged refrigerator storage is not required for this enhanced alloimmunization.

**Figure 6:**
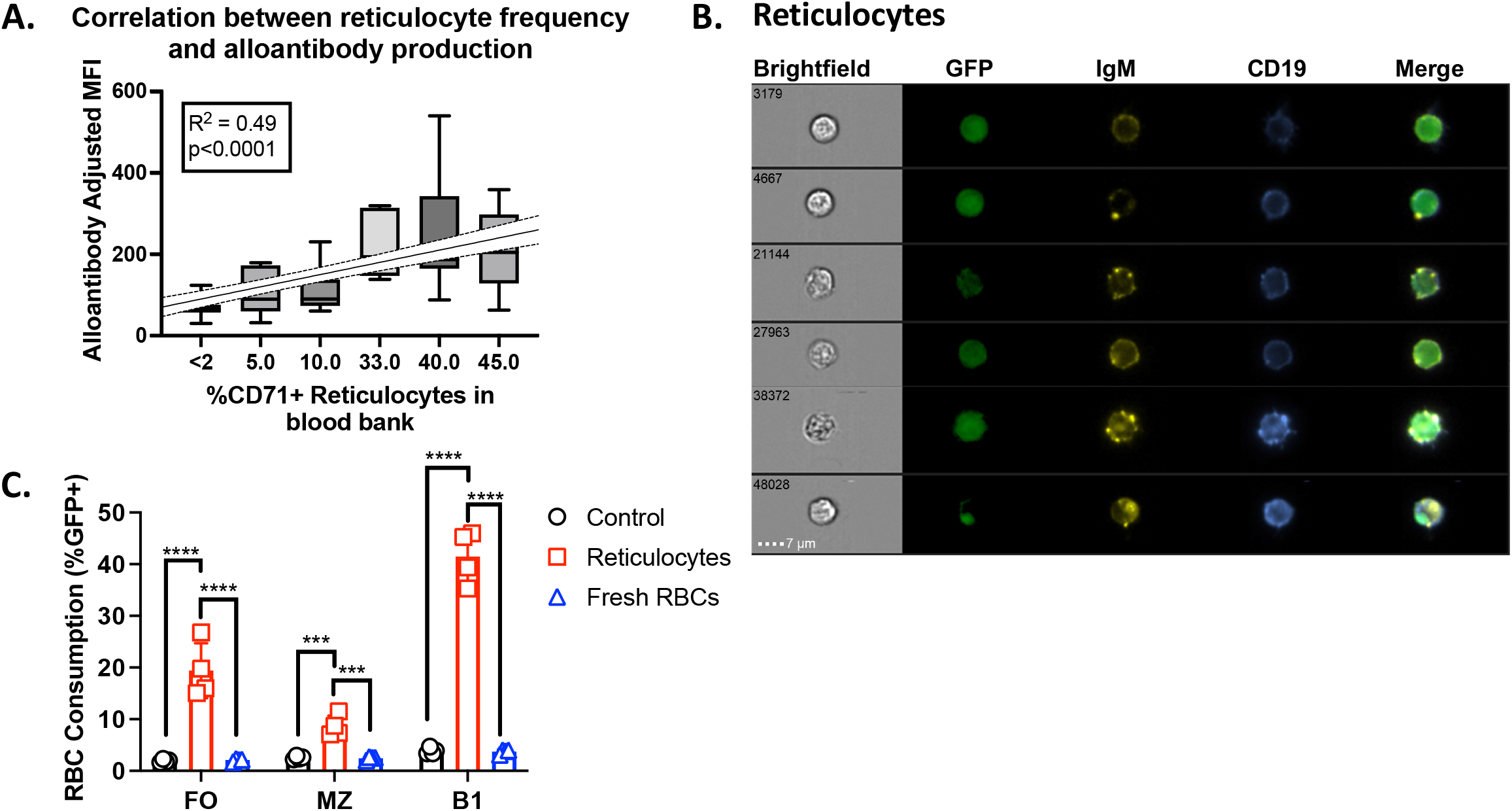
RBC alloimmunization is stimulated, even with shorter storage duration and lower reticulocyte frequencies. RBC units with defined reticulocyte frequencies were generated by titrating whole blood from phenylhydrazine (PHZ)-treated animals into RBCs collected from saline-treated animals. RBC units were leukoreduced and stored at 4°C for 1 day before transfusion into B6 recipient mice. (A) RBC alloantibodies were quantified by flow crossmatch at 14 days post-transfusion and alloantibody mean fluorescence intensity (MFI) was plotted against the RBC unit reticulocyte frequency. Data were analyzed with a one-way ANOVA and tested for linear trend. Data are cumulative of at least 3 independent experiments and each data point is one mouse, 0% (n=25), 5% (n=15), 10% (n=13), 33% (n=5), 40% (n=15), 45% (n=5). (B) Flow analysis revealed that multiple B cell subsets consumed GFP+ RBCs from reticulocyte-rich RBC units and this was confirmed visually by (C) imaging analysis. For RBC consumption, data were analyzed with a one-way ANOVA with Tukey’s multiple comparisons test; ****p<0.0001, ***p<0.001.

### TLR4 is not required for reticulocyte-mediated enhanced alloimmunization

Reticulocytes contain endogenous danger signals that can induced inflammation upon recognition by immune cells. Because stress-reticulocytes are more prone to hemolysis(20, 21) (thereby releasing free heme(35)) and contain mitochondria (enveloped in cardiolipin)(36), we hypothesized that TLR4 was required for the observed enhanced alloimmune responses upon reticulocyte-rich transfusion. To that end, B6 and TLR4-/-recipients were transfused with reticulocyte-rich or control RBC units; no significant differences in anti-HOD alloantibodies were observed 14 days post-transfusion (**Fig. 7**). Thus, these data demonstrate that reticulocyte-mediated enhanced alloimmune responses do not require TLR4 signaling. These data also rule out the potential contribution of LPS contamination in RBC unit production.

**Figure 7:**
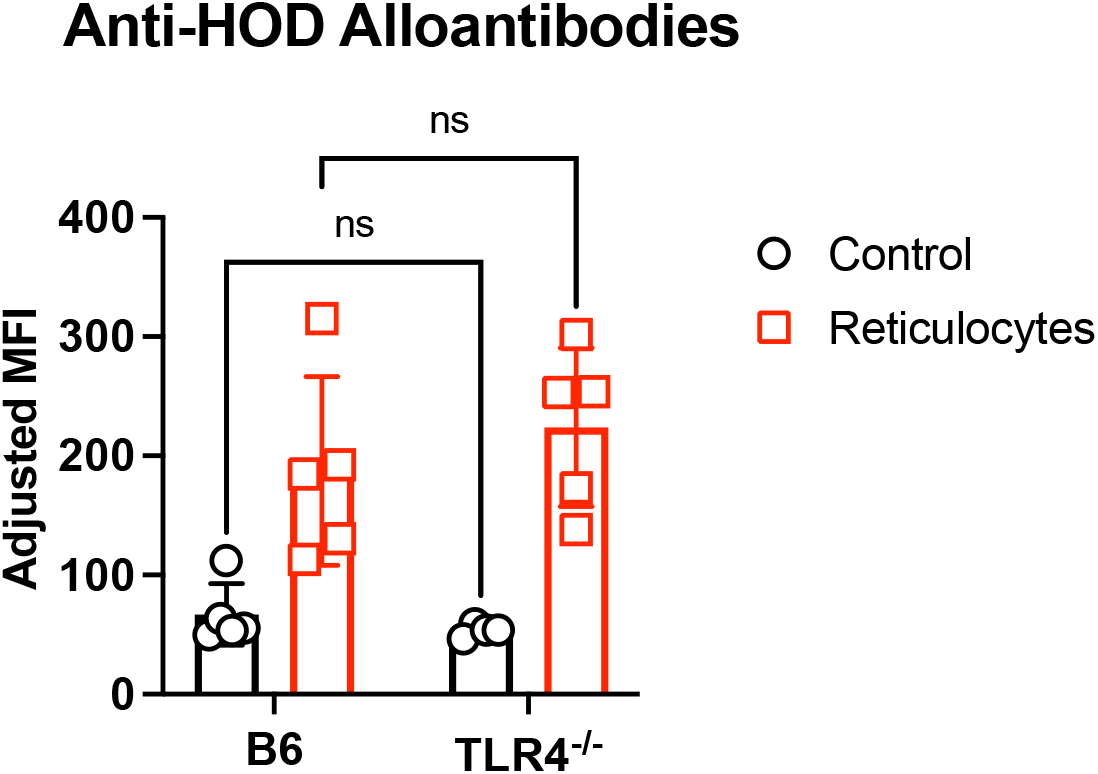
TLR4 is not required for reticulocyte-mediated enhanced RBC alloimmunization. Reticulocyte-rich and control RBC units, generated from HOD donor animals, were leukoreduced and stored 1 day at 4°C before transfusion into wild-type B6 or TLR4^-/-^ mice. Sera was collected 14 days post-transfusion and analyzed for total anti-HOD RBC alloantibodies by flow crossmatch. Data are from one experiment with 4-5 mice/group. Data were analyzed with a two-way ANOVA with Sidak’s multiple comparisons test; ns = not significant

## Discussion

These results demonstrate that increased reticulocyte count in a refrigerator-stored RBC unit can enhance transfusion-induced RBC alloimmune responses. Reticulocyte-mediated modulation of alloimmunization was dose-dependent, and was not an artifact of PHZ treatment as a similar effect was observed using reticulocytes generated from an iron deficiency/iron repletion model. Moreover, reticulocyte-enhanced alloimmunization was independent of refrigerator storage duration because transfusion of reticulocyte-rich RBC units refrigerator stored for either 1 or 6 days enhanced alloimmunization rates and alloantibody levels. To explore underlying mechanisms, transfusion of reticulocyte-rich RBC units elicited pro-inflammatory cytokines, and transfused RBCs were cleared more rapidly, preferentially by splenic B cells. Lastly, we show that the observed reticulocyte-mediated enhanced alloimmune responses do not require TLR4 signaling; these data exclude the possibility of potential LPS contamination and rule out potentially immunogenic reticulocyte-derived ligands heme and cardiolipin. Together, these findings imply that reticulocytes in RBC units may be immunogenic and could enhance alloimmune responses, even in otherwise stable recipients; this finding may have implications for transfusions into all human patients, particularly those with the highest risk for alloimmunization.

Approximately 70% of the blood supply in the USA derives from repeat volunteer blood donors(37, 38). Because blood donation typically removes ∼10% of the total blood volume, compensatory stress erythropoiesis increases production of erythroid precursors and the premature release of reticulocytes; although the minimum inter-donation interval for blood donation in the US is 56 days, elevated reticulocyte counts (over baseline) are observed even after ∼160 days after donation(15, 39). Thus, frequent donation by repeat blood donors can yield RBC units with higher reticulocyte counts. Additionally, current guidance for iron-deficient volunteers is to take iron supplements before donation, which also promotes stress erythropoiesis(15). Reticulocytes differ from their mature counterparts by: (1) containing immune-enhancing organelles (e.g., mitochondria), lipids, and residual RNA; (2) expressing surface antigens that promote adherence (e.g., CD44, CD36); and (3) being larger and less deformable. Compared to steady-state reticulocytes, stress erythropoiesis-induced reticulocytes also contain significantly more organelles (e.g., mitochondria, ribosomes), express higher levels of surface molecules mediating adherence, and have shorter circulatory lifespans due to their size, decreased deformability, and susceptibility to shear stress(18-23). Thus, reticulocytes, and especially stress-induced reticulocytes, contain and/or express potentially immunogenic components.

Recipient inflammation is associated with higher alloimmunization rates and alloantibody levels(4, 5). Because RBC alloimmunization is correlated with type I IFN levels, and because stress-induced reticulocytes contain ligands that could stimulate type I IFN production (e.g., mitochondrial DNA(13, 14)), we measured plasma IFN-α and IFN-β. Unexpectedly, no significant levels were detectable. This could be due to timing, as plasma was collected 2 hours post-transfusion, or location (e.g., production only in the spleen); additionally, it could indicate that type I IFNs are not required for reticulocyte-enhanced alloimmunization. Nonetheless, transfusion of reticulocyte-rich units elicited high levels of proinflammatory cytokines and chemokines. Other potentially immunogenic components of reticulocytes include mitochondria-derived cardiolipin and heme, both of which are TLR4 ligands. However, transfusing reticulocyte-rich RBC units into TLR4^-/-^ and control recipients led to similar alloantibody levels (**Fig. 7**), demonstrating TLR4 is dispensable for the enhanced alloimmune responses. Future studies will identify which signaling pathways are required for reticulocyte-mediated enhanced alloimmunity.

In a departure from typical RBC clearance patterns(8), B cells preferentially consumed RBCs derived from reticulocyte-rich units. Macrophages are typically responsible for steady-state removal of senescent and/or damaged RBCs(8, 34). In murine models of RBC alloimmunization, with transfusion of stored RBCs or in the context of inflammation, macrophages and DC subsets (CD11b+ and CD8+) play critical roles in initiating alloimmune responses(9, 34). Although B cell consumption of transfused RBCs is, indeed, a prerequisite for alloantibody production, very few B cells are typically observed in this process. In contrast, B cells play a major role clearing transfused RBCs from reticulocyte-rich units. Because of the high frequency of follicular and marginal zone B cells co-localizing with RBCs, it is unlikely that RBC phagocytosis is B cell receptor (BCR)-mediated; however, B1 B cells express polyreactive BCRs that bind many autoantigens, including phosphatidylserine (PS), which is expressed at high levels on stress-induced reticulocytes(40, 41). Additionally, innate B1 B cells can participate in BCR-independent endocytosis(42), although the underlying mechanisms are not yet defined. As all B cells express complement receptors and Fc receptors(43, 44), these pathways may be involved in erythrophagocytosis. Elucidating the key pathways required for B cell consumption of RBCs in this setting will be a focus of future experiments.

As anemia of many etiologies induce stress erythropoiesis, one potential limitation of these studies is that various stresses may produce differences in the resulting reticulocytes. Thus, it is possible that reticulocyte-containing RBC units generated from repeatedly phlebotomized animals may not enhance RBC alloimmunization. To address this concern, the current studies utilized two different models of anemia— chemically-induced hemolytic anemia and iron deficiency anemia followed by iron repletion—to induce reticulocytosis. Transfusion of RBC units generated with both models enhanced alloimmunization rates and alloantibody levels, as compared to controls. Another limitation is the inability to distinguish steady-state and stress-induced reticulocytes to assess their relative contributions to RBC alloimmunization; we are optimizing approaches to define a distinct phenotype for stress-induced reticulocytes, which would be essential for screening RBC donor units and attributing specific functional outcomes in response to post-transfusion clearance. Although most studies presented herein utilize artificially high reticulocyte frequencies in donor RBC units for initial phenomenological studies, clinically relevant enhanced alloimmune responses were observed with as few as 5% reticulocytes (**Fig. 6A**); it is noteworthy that the average reticulocyte frequency in unmanipulated animals in our studies is ∼2% (**Fig. 6A**), which is approximately twice the average observed in humans; this is likely due to the shortened lifespan of RBCs in a mouse (∼45 days) compared to humans (∼100-120 days)(45). Thus, while these findings may be clinically relevant to human transfusion, the study of reticulocyte-directed immune responses is biologically relevant to many other systems (e.g., neocytolysis(46), malaria(47), production of *in vitro* universal donor RBCs(48) etc.).

Transfusion of RBCs is an essential therapy in numerous diseases and is the most common inpatient therapeutic procedure requiring consent, with ∼11 million transfusions per year in the United States(49). Although RBC transfusions can be lifesaving, some patients develop alloantibodies against donor RBC blood group antigens. RBC alloimmunization is clinically significant because it can cause adverse events (e.g., hyperhemolysis) and pose barriers to future care (e.g., organ transplants). In addition, alloantibody prevention and detection, and mitigation of adverse events, requires significant medical, financial, and human resources. As such, identifying risk factors that promote alloimmunization is a high priority and would allow for reallocation of resources to patients most at-risk and provide potential therapeutic interventions. Herein, we demonstrate that the reticulocyte count in donor RBC units can enhance RBC alloimmunization, suggesting that increased reticulocyte frequencies may make RBC units more immunogenic and screening for reticulocyte count and/or lengthening the time between donations for repeat blood donors may reduce alloimmunization events.

## Author Contributions

TT, EAH, SLS and KEH designed the studies and experiments. AQ, CYK, DEG, AM, MT, and FDZ collected and analyzed data from murine experiments. MD and AD performed the metabolomics and proteomics analyses. All authors were involved in the interpretation of data. TT, EAH, SLS, and KEH wrote the manuscript. All authors contributed to the manuscript and approved the submitted version.

## Acknowledgements

The authors would like to thank Michael Kissner, Director of the Columbia Stem Cell Initiative Flow Cytometry Core, for assistance in experimental design and image acquisition with the ImageStreamX MkII imaging cytometer. Research reported in this publication using the ImageStreamX MkII imaging cytometer was performed in the Columbia University Stem Cell Initiative Flow Cytometry core facility at Columbia University Irving Medical Center and was supported by the Office of the Director, National Institutes of Health under Award Number S10OD026845. The content is solely the responsibility of the authors and does not necessarily represent the official views of the National Institutes of Health.

## References

1. Tormey CA, Hendrickson JE. Transfusion-related red blood cell alloantibodies: induction and consequences. Blood. 2019;133(17):1821–1830. doi:10.1182/blood-2018-08-833962.

2. Roubinian NH, Reese SE, Qiao H, Plimier C, Fang F, Page GP, Cable RG, Custer B, Gladwin MT, Goel R, Harris B, Hendrickson JE, Kanias T, Kleinman S, Mast AE, Sloan SR, Spencer BR, Spitalnik SL, Busch MP, Hod EA. Donor genetic and nongenetic factors affecting red blood cell transfusion effectiveness. JCI Insight. 2022 01/11/;7(1). doi:10.1172/jci.insight.152598.

3. Rapido F, Brittenham GM, Bandyopadhyay S, La Carpia F, L’Acqua C, McMahon DJ, Rebbaa A, Wojczyk BS, Netterwald J, Wang H, Schwartz J, Eisenberger A, Soffing M, Yeh R, Divgi C, Ginzburg YZ, Shaz BH, Sheth S, Francis RO, Spitalnik SL, Hod EA. Prolonged red cell storage before transfusion increases extravascular hemolysis. J Clin Invest. 2017 01/03/;127(1):375–382. doi:10.1172/JCI90837.

4. Ryder AB, Zimring JC, Hendrickson JE. Factors Influencing RBC Alloimmunization: Lessons Learned from Murine Models. Transfusion Medicine and Hemotherapy. 2014;41(6):406–419. doi:10.1159/000368995.

5. Evers D, van der Bom JG, Tijmensen J, Middelburg RA, de Haas M, Zalpuri S, de Vooght KMK, van de Kerkhof D, Visser O, Péquériaux NCV, Hudig F, Zwaginga JJ. Red cell alloimmunisation in patients with different types of infections [https://doi.org/10.1111/bjh.14307]. British Journal of Haematology. 2016 2016/12/01;175(5):956–966. doi:https://doi.org/10.1111/bjh.14307.

6. Gibb DR, Liu J, Natarajan P, Santhanakrishnan M, Madrid DJ, Eisenbarth SC, Zimring JC, Iwasaki A, Hendrickson JE. Type I IFN Is Necessary and Sufficient for Inflammation-Induced Red Blood Cell Alloimmunization in Mice. The Journal of Immunology. 2017:ji1700401. doi:10.4049/jimmunol.1700401.

7. Lee JY, Madany E, El Kadi N, Pandya S, Ng K, Yamashita M, Jefferies CA, Gibb DR. Type 1 Interferon Gene Signature Promotes RBC Alloimmunization in a Lupus Mouse Model [Original Research]. Frontiers in Immunology. 2020;11.

8. Richards AL, Hendrickson JE, Zimring JC, Hudson KE. Erythrophagocytosis by plasmacytoid dendritic cells and monocytes is enhanced during inflammation. Transfusion. 2016 2016/04/01;56(4):905–916. doi:10.1111/trf.13497.

9. Richards AL, Sheldon K, Wu X, Gruber DR, Hudson KE. The Role of the Immunological Synapse in Differential Effects of APC Subsets in Alloimmunization to Fresh, Non-stored RBCs [10.3389/fimmu.2018.02200]. Frontiers in Immunology. 2018;9:2200.

10. Rönnblom L, Eloranta M-L. The interferon signature in autoimmune diseases. Current Opinion in Rheumatology. 2013;25(2).

11. Kim M, Hwang S, Park K, Kim SY, Lee YK, Lee DS. Increased Expression of Interferon Signaling Genes in the Bone Marrow Microenvironment of Myelodysplastic Syndromes. PLoS One. 2015;10(3):e0120602. doi:10.1371/journal.pone.0120602.

12. Madany E, Lee J, Halprin C, Seo J, Baca N, Majlessipour F, Hendrickson JE, Pepkowitz SH, Hayes C, Klapper E, Gibb DR. Altered type 1 interferon responses in alloimmunized and nonalloimmunized patients with sickle cell disease. EJHaem. 2021;2(4):700–710. eng. doi:10.1002/jha2.270.

13. Moriconi C, Dzieciatkowska M, Roy M, D’Alessandro A, Roingeard P, Lee JY, Gibb DR, Tredicine M, McGill MA, Qiu A, La Carpia F, Francis RO, Hod EA, Thomas T, Picard M, Akpan IJ, Luckey CJ, Zimring JC, Spitalnik SL, Hudson K. Retention of functional mitochondria in mature red blood cells from patients with sickle cell disease.. British Journal of Haematology. 2022;198(3):574–586. eng. doi:10.1111/bjh.18287.

14. Caielli S, Cardenas J, de Jesus AA, Baisch J, Walters L, Blanck JP, Balasubramanian P, Stagnar C, Ohouo M, Hong S, Nassi L, Stewart K, Fuller J, Gu J, Banchereau JF, Wright T, Goldbach-Mansky R, Pascual V. Erythroid mitochondrial retention triggers myeloid-dependent type I interferon in human SLE. Cell. 2021 2021/08/19/;184(17):4464-4479.e19. doi:https://doi.org/10.1016/j.cell.2021.07.021.

15. Mast AE, Szabo A, Stone M, Cable RG, Spencer BR, Kiss JE. The benefits of iron supplementation following blood donation vary with baseline iron status. American Journal of Hematology. 2020;95(7):784–791. eng.

16. Wong E, Rose M, Berliner N. Disorders of Red Blood Cells. Cecil Essentials of Medicine, Tenth Edition. Vol 48. Elsevier; 2022. Chapter Disorders of Red Blood Cells; p. 489–500.

17. Stryckmans PA, Cronkite EP, Giacomelli G, Schiffer LM, Schnappauf H. The Maturation and Fate of Reticulocytes after In Vitro Labeling with Tritiated Amino Acids. Blood. 1968 1968/01/01/;31(1):33–43. doi:https://doi.org/10.1182/blood.V31.1.33.33.

18. Come SE, Shohet SB, Robinson SH. Surface Remodelling of Reticulocytes produced in Response to Erythroid Stress. Nature New Biology. 1972 1972/04/01;236(66):157–158. doi:10.1038/newbio236157a0.

19. Come SE, Shohet SB, Robinson SH. Surface Remodeling vs. Whole-Cell Hemolysis of Reticulocytes Produced With Erythroid Stimulation or Iron Deficiency Anemia. Blood. 1974 1974/12/01/;44(6):817–830. doi:https://doi.org/10.1182/blood.V44.6.817.817.

20. Noble NA, Xu Q-P, Hoge LL. Reticulocytes II: Reexamination of the In Vivo Survival of Stress Reticulocytes. Blood. 1990 1990/05/01/;75(9):1877–1882. doi:https://doi.org/10.1182/blood.V75.9.1877.1877.

21. Robinson SH, Tsong M. Hemolysis of “stress” reticulocytes: a source of erythropoietic bilirubin formation. J Clin Invest. 1970 05/01/;49(5):1025–1034. doi:10.1172/JCI106302.

22. Carden MA, Fasano RM, Meier ER. Not all red cells sickle the same: Contributions of the reticulocyte to disease pathology in sickle cell anemia. Blood Reviews. 2020 2020/03/01/;40:100637. doi:https://doi.org/10.1016/j.blre.2019.100637.

23. Sawadogo D, Tolo-Dilkébié A, Sangaré M, Aguéhoundé N, Kassi H, Latte T. Influence of the Clinical Status on Stress Reticulocytes, CD36 and CD49d of SSFA 2 Homozygous Sickle Cell Patients Followed in Abidjan. Advances in Hematology. 2014 2014/02/27;2014:273860. doi:10.1155/2014/273860.

24. Desmarets M, Cadwell CM, Peterson KR, Neades R, Zimring JC. Minor histocompatibility antigens on transfused leukoreduced units of red blood cells induce bone marrow transplant rejection in a mouse model. Blood. 2009;114(11):2315–2322. eng. Epub 2009/06/12. doi:10.1182/blood-2009-04-214387. Cited in: Pubmed; PMID 19525479.

25. Bandyopadhyay S, Brittenham GM, Francis RO, Zimring JC, Hod EA, Spitalnik SL. Iron-deficient erythropoiesis in blood donors and red blood cell recovery after transfusion: initial studies with a mouse model. Blood Transfusion. 2017;15(2):158–164. eng. doi:10.2450/2017.0349-16.

26. Kim CY, Johnson H, Peltier S, Spitalnik SL, Hod EA, Francis RO, Hudson KE, Stone EF, Gordy DE, Fu X, Zimring JC, Amireault P, Buehler PW, Wilson RB, D’Alessandro A, Shchepinov MS, Thomas T. Deuterated Linoleic Acid Attenuates the RBC Storage Lesion in a Mouse Model of Poor RBC Storage [Original Research]. Frontiers in Physiology. 2022;13.

27. Qiu A, Miller A, Zotti FD, Santhanakrishnan M, Hendrickson JE, Tredicine M, Stowell SR, Luckey CJ, Zimring JC, Hudson KE. FcγRIV is required for IgG2c mediated enhancement of RBC alloimmunization [Original Research]. Frontiers in Immunology. 2022;13.

28. Richards AL, Qiu A, Zotti FD, Sheldon K, Usaneerungrueng C, Gruber DR, Hudson KE. Autoantigen presentation by splenic dendritic cells is required for RBC-specific autoimmunity [https://doi.org/10.1111/trf.16191]. Transfusion. 2021 2021/01/01;61(1):225–235. doi:https://doi.org/10.1111/trf.16191.

29. Richards AL, Howie HL, Kapp LM, Hendrickson JE, Zimring JC, Hudson KE. Innate B-1 B Cells Are Not Enriched in Red Blood Cell Autoimmune Mice: Importance of B Cell Receptor Transgenic Selection [10.3389/fimmu.2017.01366]. Frontiers in Immunology. 2017;8:1366.

30. Nemkov T, Reisz JA, Gehrke S, Hansen KC, D’Alessandro A. High-Throughput Metabolomics: Isocratic and Gradient Mass Spectrometry-Based Methods. Methods Mol Biol. 2019;1978:13–26. eng.

31. Pang Z, Chong J, Zhou G, de Lima Morais DA, Chang L, Barrette M, Gauthier C, Jacques P-É, Li S, Xia J. MetaboAnalyst 5.0: narrowing the gap between raw spectra and functional insights. Nucleic Acids Research. 2021;49(W1):W388–W396. doi:10.1093/nar/gkab382.

32. Thomas T, Stefanoni D, Dzieciatkowska M, Issaian A, Nemkov T, Hill RC, Francis RO, Hudson KE, Buehler PW, Zimring JC, Hod EA, Hansen KC, Spitalnik SL, D’Alessandro A. Evidence of Structural Protein Damage and Membrane Lipid Remodeling in Red Blood Cells from COVID-19 Patients. Journal of Proteome Research. 2020 2020/11/06;19(11):4455–4469. doi:10.1021/acs.jproteome.0c00606.

33. Shetlar MD, Hill HA. Reactions of hemoglobin with phenylhydrazine: a review of selected aspects. Environmental Health Perspectives. 1985 1985/12/01;64:265–281. doi:10.1289/ehp.8564265.

34. Youssef L, Rebbaa A, Pampou S, Stockwell BR, Spitalnik S. Transfusion of Storage-Damaged Red Blood Cells Induces Ferroptosis in Splenic Macrophages. Blood. 2017 2017/12/08/;130:3567. doi:https://doi.org/10.1182/blood.V130.Suppl_1.3567.3567.

35. Janciauskiene S, Vijayan V, Immenschuh S. TLR4 Signaling by Heme and the Role of Heme-Binding Blood Proteins [Review]. Frontiers in Immunology. 2020;11.

36. Pizzuto M, Pelegrin P. Cardiolipin in Immune Signaling and Cell Death. Trends in Cell Biology. 2020;30(11):892–903. doi:10.1016/j.tcb.2020.09.004.

37. Whitaker B, Rajbhandary S, Kleinman S, Harris A, Kamani N. Trends in United States blood collection and transfusion: results from the 2013 AABB Blood Collection, Utilization, and Patient Blood Management Survey [https://doi.org/10.1111/trf.13676]. Transfusion. 2016 2016/09/01;56(9):2173–2183. doi:https://doi.org/10.1111/trf.13676.

38. Kiss JE, Vassallo RR. How do we manage iron deficiency after blood donation? [https://doi.org/10.1111/bjh.15136]. British Journal of Haematology. 2018 2018/06/01;181(5):590–603. doi:https://doi.org/10.1111/bjh.15136.

39. Schotten N, Pasker-de Jong PCM, Moretti D, Zimmermann MB, Geurts-Moespot AJ, Swinkels DW, van Kraaij MGJ. The donation interval of 56 days requires extension to 180 days for whole blood donors to recover from changes in iron metabolism. Blood. 2016;128(17):2185–2188. doi:10.1182/blood-2016-04-709451.

40. Hosseini H, Li Y, Kanellakis P, Tay C, Cao A, Tipping P, Bobik A, Toh B-H, Kyaw T. Phosphatidylserine liposomes mimic apoptotic cells to attenuate atherosclerosis by expanding polyreactive IgM producing B1a lymphocytes. Cardiovascular Research. 2015;106(3):443–452. doi:10.1093/cvr/cvv037.

41. Baumgarth N. How specific is too specific? B-cell responses to viral infections reveal the importance of breadth over depth [https://doi.org/10.1111/imr.12094]. Immunological Reviews. 2013 2013/09/01;255(1):82–94. doi:https://doi.org/10.1111/imr.12094.

42. Parra D, Rieger AM, Li J, Zhang Y-A, Randall LM, Hunter CA, Barreda DR, Sunyer JO. Pivotal Advance: Peritoneal cavity B-1 B cells have phagocytic and microbicidal capacities and present phagocytosed antigen to CD4+ T cells [https://doi.org/10.1189/jlb.0711372]. Journal of Leukocyte Biology. 2012 2012/04/01;91(4):525–536. doi:https://doi.org/10.1189/jlb.0711372.

43. Liu J, Wang Y, Xiong E, Hong R, Lu Q, Ohno H, Wang J-Y. Role of the IgM Fc Receptor in Immunity and Tolerance [Review]. Frontiers in Immunology. 2019;10.

44. Dunkelberger JR, Song W-C. Complement and its role in innate and adaptive immune responses. Cell Research. 2010 2010/01/01;20(1):34–50. doi:10.1038/cr.2009.139.

45. Krzyzanski W, Brier ME, Creed TM, Gaweda AE. Reticulocyte-based estimation of red blood cell lifespan. Experimental Hematology. 2013;41(9):817–822. doi:10.1016/j.exphem.2013.05.001.

46. Mairbäurl H. Neocytolysis: How to Get Rid of the Extra Erythrocytes Formed by Stress Erythropoiesis Upon Descent From High Altitude [Mini Review]. Frontiers in Physiology. 2018;9.

47. Lim C, Pereira L, Saliba KS, Mascarenhas A, Maki JN, Chery L, Gomes E, Rathod PK, Duraisingh MT. Reticulocyte Preference and Stage Development of Plasmodium vivax Isolates. The Journal of Infectious Diseases. 2016;214(7):1081–1084. doi:10.1093/infdis/jiw303.

48. Pellegrin S, Severn CE, Toye AM. Towards manufactured red blood cells for the treatment of inherited anemia. Haematologica. 2021 05/27;106(9):2304–2311. doi:10.3324/haematol.2020.268847.

49. Jones JM, Sapiano MRP, Mowla S, Bota D, Berger JJ, Basavaraju SV. Has the trend of declining blood transfusions in the United States ended? Findings of the 2019 National Blood Collection and Utilization Survey. Transfusion. 2021;61;Suppl 2(1537-2995 (Electronic)):S1–S10. eng.

