## Supplemental Data for "Reticulocytes in donor RBC units enhance RBC alloimmunization"

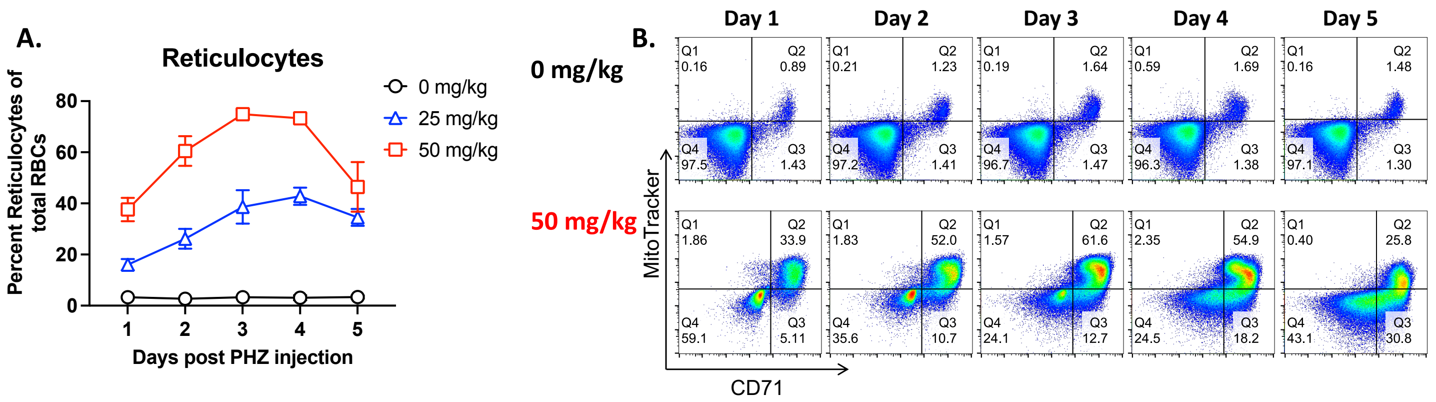


**Supplemental Figure 1: Phenylhydrazine (PHZ) dose titration.** B6 mice received 2 i.p. injections of (1) saline as a control (0 mg/kg; black circle), or (2) 25 mg/kg PHZ (blue triangle), or 3) 50mg/kg PHZ (red square). Peripheral blood was collected daily for 6 days and stained to detect CD71+ reticulocytes, CD71- mature RBCs, and mitochondria. Cells were gated on CD45-CD41- singlets to exclude white blood cells and platelets. (A) The percentages of CD71+ reticulocytes out of total CD41-CD45- RBCs were calculated; data are mean ± SD. Cell-permeable MitoTracker dye identified reticulocytes with detectable mitochondria; (B) representative flow plots of CD71 and MitoTracker staining after 50 mg/kg PHZ or PBS (0 mg/kg) treatment; mitochondria-positive reticulocytes are in Q2. Data are representative of 2 independent experiments with 5 mice/group.

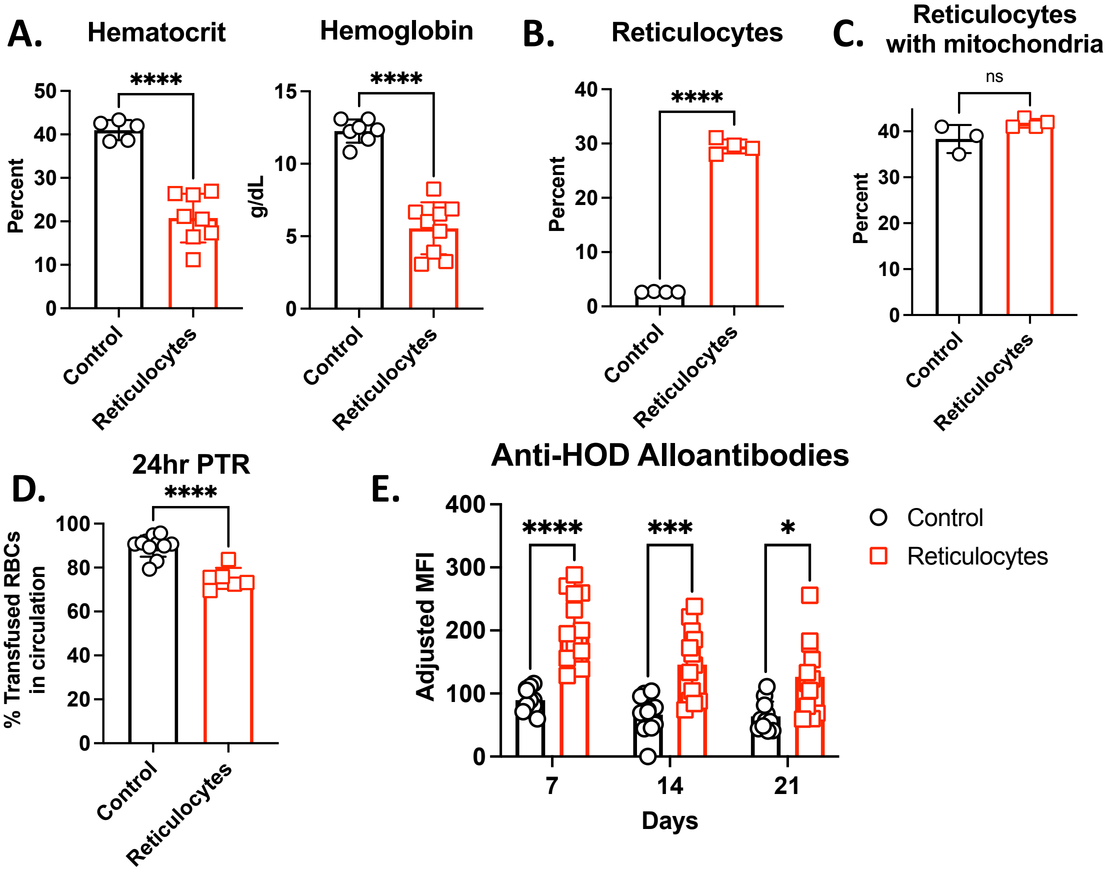


**Supplemental Figure 2: Transfusion of refrigerator-stored reticulocyte-rich RBC units from an iron-deficient mouse model led to increased RBC alloantibodies.** Weanling Hen egg lysozyme-Ovalbumin-Duffy (HOD) mice were placed on an iron-deficient (referred to as “Reticulocytes”) or an iron-replete diet (referred to as “Control”). After 4 weeks on iron-defined diets, (A) hematocrit and hemoglobin were determined to assess anemia. All mice then received 5mg of iron dextran, and whole blood was collected 4 days later into 14% CPDA-1 by cardiac puncture. The frequencies of (B) total reticulocytes and (C) the percent of mitochondria-positive reticulocytes was calculated. RBC units were refrigerator-stored for 6 days before transfusion into B6 recipients. (D) Post-transfusion recovery (PTR) was determined. (E) Sera, collected weekly, were analyzed for anti-HOD alloantibodies by flow crossmatch. The experiment was completed once, with 11-13 transfusion recipients/group. Data were analyzed with an unpaired t-test for 2 groups or one way ANOVA with Sidak’s multiple comparisons test; ****$p<0.0001$, ***$p<0.001$, *$p<0.05$, ns = not significant

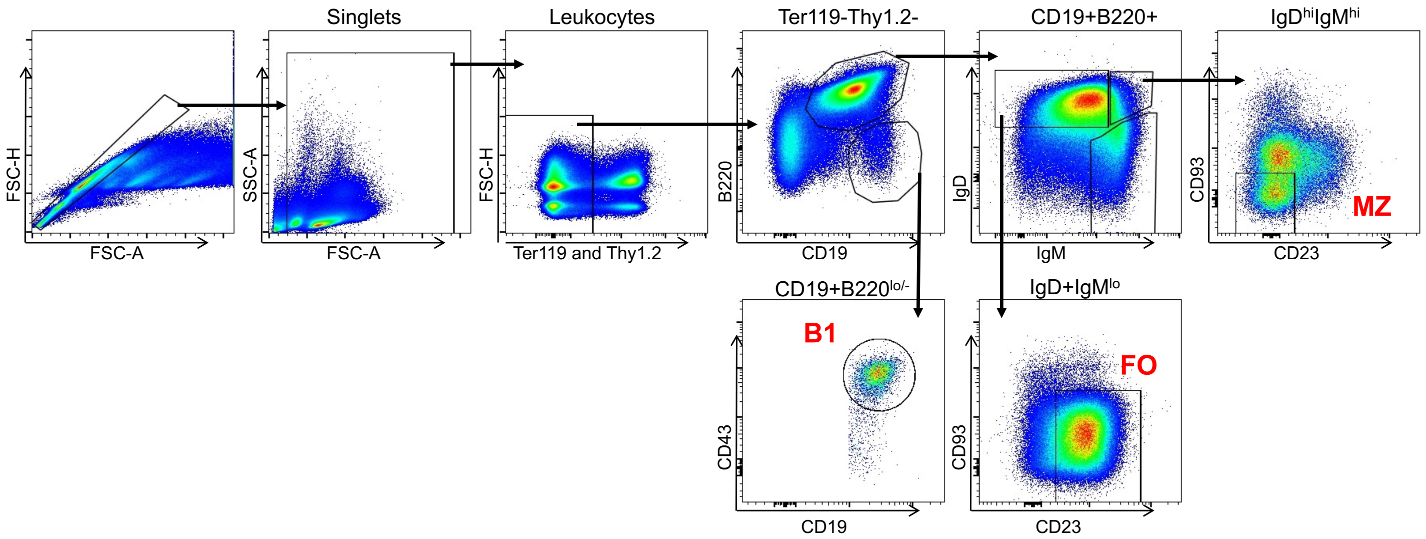


**Supplemental Figure 3: B cell gating strategy**

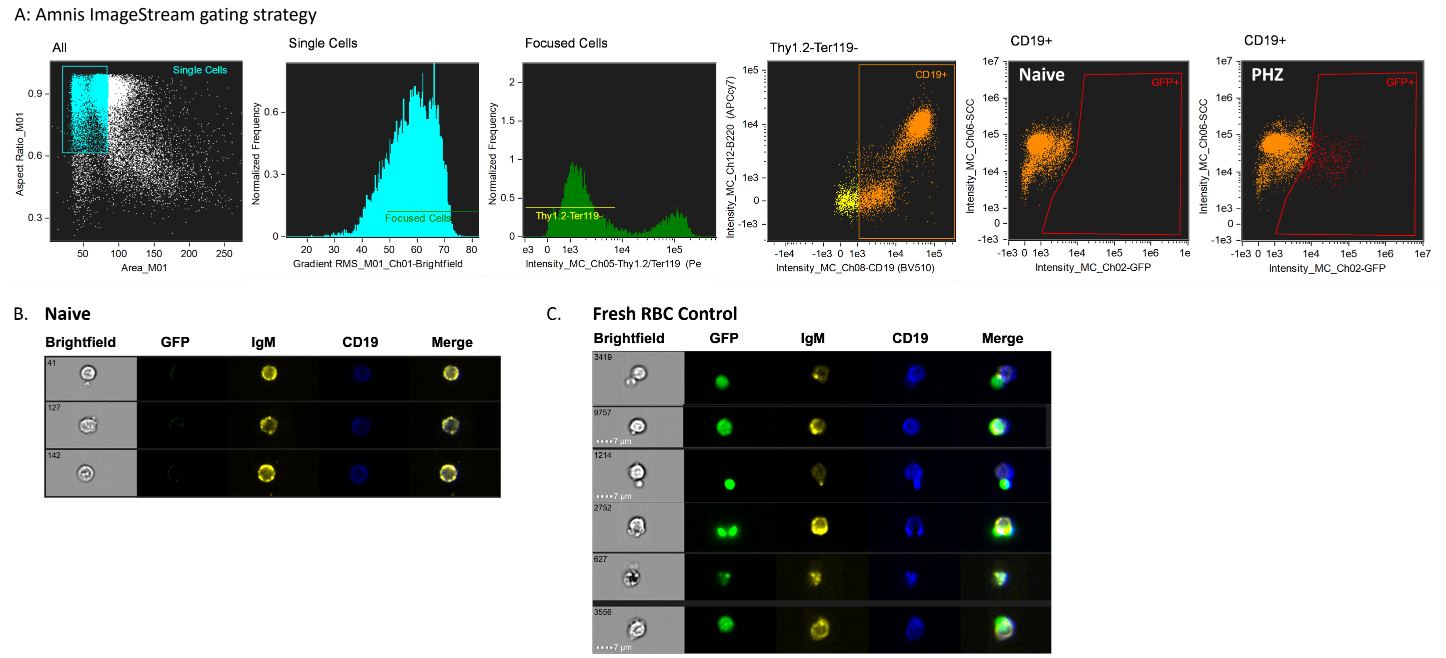


**Supplemental Figure 4: Amnis ImageStream gating strategy and GFP+ B cells from mice transfused with fresh GFP RBCs.** (A) Gating strategy for Amnis image analysis. (B) GFP gating and background signal were determined on B cells from naïve, non-transfused animals. (C) Images from GFP+ B cells collected from recipient mice transfused with fresh GFP RBCs.

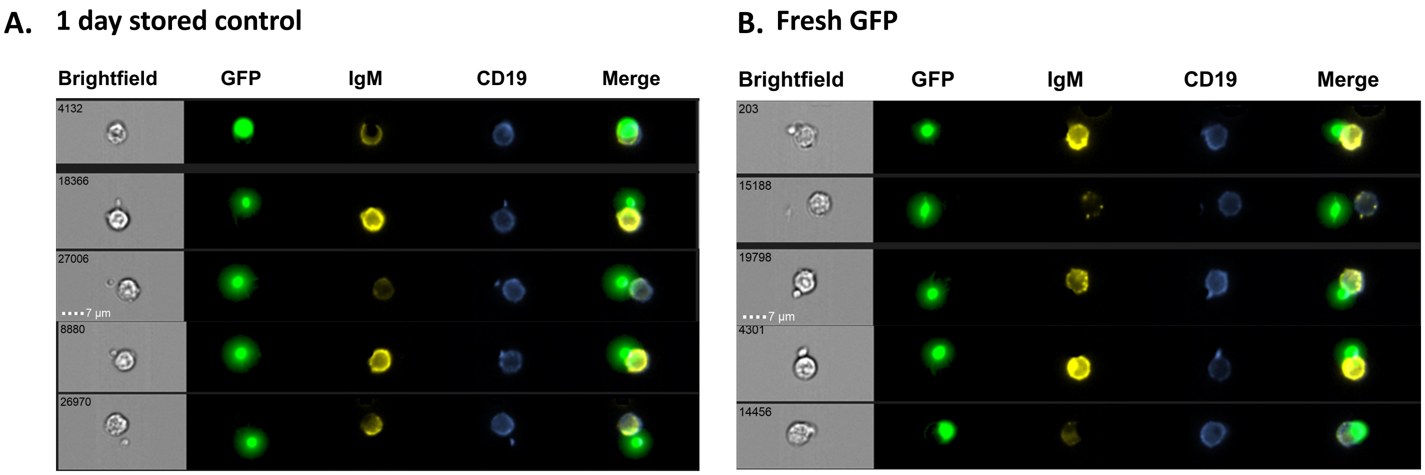


**Supplemental Figure 5: Images of GFP+ B cells from recipients transfused with 1 day stored control (A) or fresh GFP (B) RBC units.**
